# *Saccharomyces cerevisiae* histone Chaperones FACT and Spt6 modulate Pol II and histone occupancy genome-wide

**DOI:** 10.1101/265074

**Authors:** Rakesh Pathak, Priyanka Singh, Sudha Ananthakrishnan, Sarah Adamczyk, Olivia Schimmel, Chhabi K. Govind

**Author notes:** Corresponding author: Chhabi K. Govind, Department of Biological Sciences, Mathematics and Science Center (MSC), Room number 333, Oakland University, Rochester, MI-48085, USA.

## Abstract

Histone chaperones, chromatin remodelers, and histone modifying complexes play a critical role in alleviating the nucleosomal barrier. Here, we have examined the role of two highly conserved yeast (*Saccharomyces cerevisiae*) histone chaperones, FACT and Spt6, in regulating transcription and histone occupancy. We show that the H3 tail contributes to the recruitment of FACT to coding sequences in a manner dependent on acetylation. We found that deleting a H3 HAT Gcn5 or mutating lysines on the H3 tail impairs FACT recruitment at *ADH1* and *ARG1* genes. However, deleting the H4 tail or mutating the H4 lysines failed to dampen FACT occupancy in coding regions. Additionally, we show that FACT-depletion greatly reduces Pol II occupancy in the 5’ ends genome-wide. By contrast, Spt6-depletion led to reduction in Pol II occupancy towards the 3’ end, in a manner dependent on the gene-length. Severe transcription and histone eviction defects were also observed in a strain that was impaired for Spt6 recruitment (*spt6*Δ202) and depleted of FACT. Importantly, the severity of the defect strongly correlated with WT Pol II occupancies at these genes, indicating critical roles of Spt6 and Spt16 in promoting high-level transcription. Collectively, our study shows cooperation, as well as redundancy between chaperones, FACT and Spt6, in regulating transcription and chromatin in coding regions of transcribed genes.

## INTRODUCTION

The nucleosome is the fundamental unit of chromatin and is composed of ~ 147 bp of DNA wrapped around a histone octamer consisting of two copies of histones H2A, H2B, H3 and H4. Nucleosomes pose a significant impediment to all steps of transcription, including the steps of initiation and elongation of RNA polymerase II (Pol II) through the coding sequences (CDS) (Li *et al.* 2007). There are two principle mechanisms suggested to alleviate nucleosomal barrier: I) removal of the H2A-H2B dimer to generate hexamers that can be readily overcome by elongating polymerases (Kireeva *et al.* 2002), and II) complete removal of histone octamers leading to reduced nucleosomal density across the transcribing genes (Lee *et al.* 2004; Dion *et al.* 2007).

Various factors and enzyme complexes have been implicated in removing the nucleosomal barriers *in vivo* (Li *et al.* 2007). For example, acetylation of histone tails facilitates histone eviction by weakening histone-DNA interactions (Govind *et al.* 2007; Govind *et al.* 2010) and by promoting recruitment of ATP-dependent remodelers, such as RSC and SWI/SNF (Hassan *et al.* 2001; Dechassa *et al.* 2010; Spain *et al.* 2014). While nucleosome disassembly is important for transcription, it is also important to restore the chromatin structure in the wake of transcription. Histone chaperones, many of which are cotranscriptionally recruited to actively transcribing genes, are implicated in performing this function (Gurard-Levin *et al.* 2014).

Two such chaperones, the FACT complex (Spt16/SSRP1 in humans and Spt16/Pob3 in yeast) and Spt6, are enriched in transcribed coding regions (Andrulis *et al.* 2000; Kaplan *et al.* 2000; Krogan *et al.* 2002; Mason and Struhl 2003; Mayer *et al.* 2010; Formosa 2012; Burugula *et al.* 2014), suggesting a role for these factors in regulating transcription and in maintaining chromatin integrity. Human FACT recognizes and displaces one of the H2A/H2B dimers from the nucleosome, and promotes transcription on a chromatin template, *in vitro.* In addition, it can assemble all four histones on the DNA (Belotserkovskaya *et al.* 2003). Yeast (*S. cerevisiae*) FACT also interacts with all four histones and displays a strong affinity toward intact nucleosomes (Formosa *et al.* 2001; VanDemark *et al.* 2008). Multiple domains within the FACT subunits Spt16 and Pob3 are implicated in binding and chaperoning histones (Hondele and Ladurner 2011; Formosa 2012). The N-terminal domain (peptidase-like) of *S. pombe* Spt16, for example, binds to both core histones as well as histone N-terminal tails (Stuwe *et al.* 2008), and the M-domain of *Chaetomium thermophilum* Spt16 recognized H2A/H2B with affinity similar to that observed with full-length Spt16 (Hondele *et al.* 2013). However, in *S. cerevisiae*, the C-terminal regions of both Spt16 and Pob3 were defined as H2A/H2B binding domains (VanDemark *et al.* 2008; Kemble *et al.* 2015). Whether these different domains are required for interacting with nucleosomes in a context-dependent manner remains to be seen.

FACT is shown to promote reassembly of the displaced H3/H4 in the *ADH1, ADH2* and *STE3* coding regions (Jamai *et al.* 2009) implicating FACT in restoring chromatin behind elongating Pol II. The role of FACT (and Spt6) in regulating histone occupancy has been examined at a genome-wide scale (van Bakel *et al.* 2013; Jeronimo *et al.* 2015). Impairing FACT function leads to reduced histone occupancy and also results in aberrant incorporation of a histone variant H2AZ (Jeronimo *et al.* 2015). Additionally, recruitment of FACT to the *HO* promoter is shown to promote histone eviction (Takahata *et al.* 2009). Likewise, FACT helps in evicting H2A/H2B from the promoter of the PHO5 gene, upon induction (Ransom *et al.* 2009).

Gene-specific studies have shown a role for FACT in regulating transcription. For example, FACT mutants impaired Pol II and TBP recruitment at the *GAL1* promoter, implicating a role for FACT in regulating transcription at the initiation step (Biswas *et al.* 2006; Fleming *et al.* 2008). However, moderate reductions in Pol II occupancy were observed in the 3’ ORFs of *GAL1* and *PHO5* but not in *LacZ* or *YAT1* genes expressed on plasmids under the control of *GAL1* promoter (Jimeno-Gonzalez *et al.* 2006). Likewise, transcription defects were observed only at *ADH1* and not at *ADH2* or *STE3*, despite all three genes showing a histone reassembly defect (Jamai *et al.* 2009), making it unclear if FACT is generally required for transcription. In addition to transcription initiation, FACT is shown to participate in promoting Pol II processivity and elongation rate at *GAL1* gene, in conjunction with the H2B ubiquitination (Fleming *et al.* 2008).

FACT shares many functional similarities with another histone chaperone, known as Spt6, which interacts with H3/H4 (Kaplan *et al.* 2003; Mayer *et al.* 2010), and also with H2A/H2B (McCullough *et al.* 2015). Loss of Spt6 function leads to reduced histone occupancy over transcribed regions, suggesting a role for Spt6 in cotranscriptional histone reassembly (Ivanovska *et al.* 2011; Perales *et al.* 2013; van Bakel *et al.* 2013; Jeronimo *et al.* 2015). Spt6, along with the FACT complex, is also implicated in preventing spurious incorporation of H2AZ in coding regions (Jeronimo *et al.* 2015) and thereby helps in restricting H2AZ to promoter nucleosomes (Billon and Cote 2013). Therefore, Spt6 and FACT play important roles in maintaining chromatin integrity. Consistent with this, widespread aberrant transcription was observed in the cells deficient of Spt6 or FACT (Cheung *et al.* 2008; van Bakel *et al.* 2013).

Spt6 mutants have also been shown to alter histone modifications and reduce transcription genome-wide (Degennaro *et al.* 2013; Kato *et al.* 2013; Perales *et al.* 2013). Spt6 possesses a tandem SH2 (tSH2) domain at its C-terminus that interacts with phosphorylated Pol II CTD, *in vitro* (Dengl *et al.* 2009; Close *et al.* 2011; Liu *et al.* 2011) and this domain is implicated in promoting Spt6 recruitment to transcribed genes, *in vivo* (Mayer *et al.* 2010; Mayer *et al.* 2012; Burugula *et al.* 2014).

In this study, we show that the acetylated histone H3 tail contributes to efficient recruitment of FACT to transcribed coding sequences in *S. cerevisiae.* Depleting Spt16 subunit of FACT elicits a greater reduction in Pol II occupancies in the 5’ends of the genes, suggesting a potential role for FACT in the early elongation steps of transcription. In contrast, Pol II occupancies were reduced more towards the 3’ end in Spt6-depleted cells, suggestive of processivity defects. Together, our results suggest a spatial regulation of transcription by these two highly conserved histone chaperones. Importantly, depleting Spt16 in a strain compromised for Spt6 recruitment evoked severe defects in transcription and histone eviction, suggesting that these chaperones may cooperate to promote a high-level transcription by modulating histone occupancies across transcribed genes.

## MATERIALS AND METHODS

### Yeast strains and growth conditions

The yeast strains used in this study are listed in Table S1. The cells were grown to absorbance A_600_ of 0.5-0.6 in synthetic complete (SC) media lacking isoleucine/valine, and treated with sulfometuron methyl (SM; 0.6 µg/ml) for 30 minutes to induce Gcn4 targets. Esa1 in the *gcn5*Δ*/esa1ts* strain was inactivated by shifting the cultures grown at 25°C to 37°C for ~1.5 hours prior to induction by SM. Spt16 and Spt6 were depleted in *SPT16-TET* and *SPT6-TET* (Hughes *et al.* 2000; Mnaimneh *et al.* 2004) cells by growing these strains in the presence of 10 µg/ml doxycycline overnight, and sub-culturing the overnight cultures in 100 ml of synthetic complete (SC) media with 10 µg/ml doxycycline to an OD_600_ of 0.6.

### Coimmunoprecipitation Assay

The coimmunoprecipitation experiments were performed as described previously (Govind *et al.* 2010). The HA-tagged H2B or Spt16-Myc tagged WT and *H3*Δ*1-28* strains were resuspended in 500 µl of lysis buffer (50 mM Tris-HCl [pH-7.5], 50 mM HEPES-KOH [pH 7.9], 10 mM MgSO_4_, 100 mM (NH_4_)_2_SO_4_, 12.5 mM KOAc, 0.01% NP-40, 20% Glycerol, 1ug/ml Pepstatin A, 100 mM PMSF, 1ug/ml Leupeptin;) and 500 µl of glass beads, and disrupted by vortexing (18 seconds x 8 times, and 150 seconds on ice between each agitation cycle). Whole cell extracts were incubated overnight with magnetic beads that were pre-conjugated to anti-Myc or anti-HA antibodies in 100 µl 4X MTB buffer (200 mM HEPES-KOH [pH 7.9], 800 mM KOAc, 54 mM MgOAc2, 40% Glycerol, 0.04% NP-40, 400 mM PMSF, 4 ug/ml Pepstatin, 4ug/ml Leupeptin), and washed 5 times with the wash buffer (50 mM Tris-HCl (pH-8.0), 0.3% NP-40, 500 mM NaCl, 10% Glycerol, 1mM PMSF, 1ug/ml Leupeptin, and 1ug/ml Pepstatin). Immunoprecipitates were analyzed by western blot using the following antibodies: anti-Myc (Roche), anti-HA (Roche), anti-Spt16 and anti-Spt6 antibodies (kindly provided by Tim Formosa). The signal intensities were quantified using Image Studio lite version 5.2 (LI-COR Biosciences).

### ChIP and ChIP-chip

The cultures were crosslinked with formaldehyde and processed for chromatin immunoprecipitation as described previously (Govind *et al.* 2012). ChIPs were performed using antibodies, anti-Myc (Roche), anti-Rpb3 (Neoclone), and anti-H3 (Abcam). ChIP DNA and the related input DNA was amplified using the primers against specific regions. 5µl of ChIP dye (15% Ficol, 0.25% bromophenol blue in 1X TBE) and SYBR green dye were added in the PCR products, resolved on 8% TBE gels and visualized on a phosphorimager and quantified using ImageQuant 5.1 software. The fold enrichments were determined by taking the ratios of the ChIP signal for gene of interest, and the signal obtained for the *POL1* used as an internal control and dividing by the ratios obtained for the related input samples (ChIP/Input fold enrichment: ChIP intensities [*ARG1/POL1*]) / input intensities [*ARG1/POL1*]). The ChIP experiments were performed using at least three independent cultures, and PCR reactions were conducted at least in duplicates. The error bars represent standard error of mean (SEM).

For ChIP-chip experiments, ChIP and related input DNA samples were amplified, from at least two biological replicates, using the GenomePlex complete whole genome amplification (WGA) kit (Sigma, cat # WGA2), according to the manufacturer’s instructions. The amplified ChIP DNA and input DNA were purified by using PCR cleanup kit (Qiagen Cat # 28104) and the DNA was quantified by NanoDrop. The samples were hybridized on Agilent 4×44 arrays (G4493A) after labeling the ChIP and input DNA with Alexa555 and Alexa647 fluorescent dyes, respectively, as per the manufacturer’s instructions at genomic core facility at Michigan State University. The arrays were scanned using Agilent scanner, and data was extracted with the Feature Extraction software (Agilent) as described previously (Spain *et al.* 2014).

### Bioinformatics Analysis

The data extracted with the Feature Extraction software (Agilent) was normalized using Limma package from Bioconductor, as described previously (Venkatesh *et al.* 2012). The genes were divided into 10 equal sized bins, with the two bins assigned to the region 500 bp upstream of the transcription start site (TSS) and two bins to the 500 bp to the region downstream of the transcription end site (TES). The average probe enrichment values were assigned to the closest bin according to the probe location, and a 10 bin matrix was generated using a PERL script. Genes corresponding to the majority of dubious ORFs, tRNA genes, small nuclear RNA genes as well as autonomously replicating sequences (ARS) were removed from the dataset. The enrichments in the 6 bins between TSS and TES were averaged to obtain an average ORF occupancy. The genes for analysis were selected on the basis of ORF enrichment. The genes <500 bp in length were removed from the analyses. The versatile aggregate profiler (Brunelle *et al.* 2015) was used to generate gene-average profile. The genes were split in the middle, and the probe intensities were aligned to the TSS for the first half and to the TES for the second half of the genes.

### Box-plots

Center lines show the median and box limits indicate the 25th and 75th percentiles as determined by R software. Whiskers extend 1.5 times the interquartile range from the 25th and 75th percentiles and outliers are represented by dots.

### Data Availability

Strains generated in this study, and the sequences of primers used for ChIP analysis are available upon request. The Gene Expression Omnibus accession number for the ChIP-chip data reported in this paper is GSE69642.

## RESULTS

### Histone H3 N-terminal tails facilitate FACT interaction with chromatin *in vivo*

While Spt16 is enriched in coding regions of actively transcribing genes (Mason and Struhl 2003; Mayer *et al.* 2010), the mechanisms by which it is recruited remains to be established. Given that the FACT complex interacts with nucleosomes *in vitro* (Belotserkovskaya *et al.* 2003), it can be recruited through nucleosome interactions. Moreover, Spt16 interaction was greatly reduced with the nucleosomes lacking the histone N-terminal tails (NTTs) implicating histone tails in promoting Spt16-nucleosome interactions (Stuwe *et al.* 2008; VanDemark *et al.* 2008). Significantly, removal of the H3 and H4 tails, but not of the H2A-H2B tails, abolished Spt16-histone interactions *in vitro* (Winkler *et al.* 2011). The H3 or H4 tails therefore could facilitate recruitment/retention of the FACT complex. To test this possibility, we examined yeast Spt16 occupancy by ChIP in histone mutants lacking the H3 or H4 N-terminal tail. Spt16 occupancy in the *ARG1* 5’ and 3’ ORFs was reduced by ~50% in the H3 mutants lacking 1-20 (*H3*Δ*1-20*) or 1-28 (*H3*Δ*1-28*) N-terminal residues (Figure 1A). In comparison, in the H4 tail mutant (*H4*Δ*1-16)*, only a small (~20%) reduction was observed in the *ARG1* 3’ ORF (Figure 1A). A substantial reduction in Spt16 occupancy (~80%) was also observed at a constitutively expressed *ADH1* gene in the H3 mutants (Figure 1B), but not in the H4 mutant (Figure S1A). Although, FACT/Spt16 interacts with both H3 and H4 tail peptides, *in vitro*, (Stuwe *et al.* 2008; VanDemark *et al.* 2008) it appears that the H3 tail may help in recruiting or retaining FACT to its target genes, *in vivo.* Consistent with this idea, reduced occupancy of Spt16 was also observed in the coding regions of *PYK1, PMA1* and *GLY1* genes (Figure 1C). In contrast to the impaired Spt16 occupancy in the H3 mutant, for most genes, Pol II occupancies were comparable in WT and the H3 mutants, except for the *GLY1* gene, which showed a small reduction in Pol II occupancy (Figure S1B).

**Figure 1:**
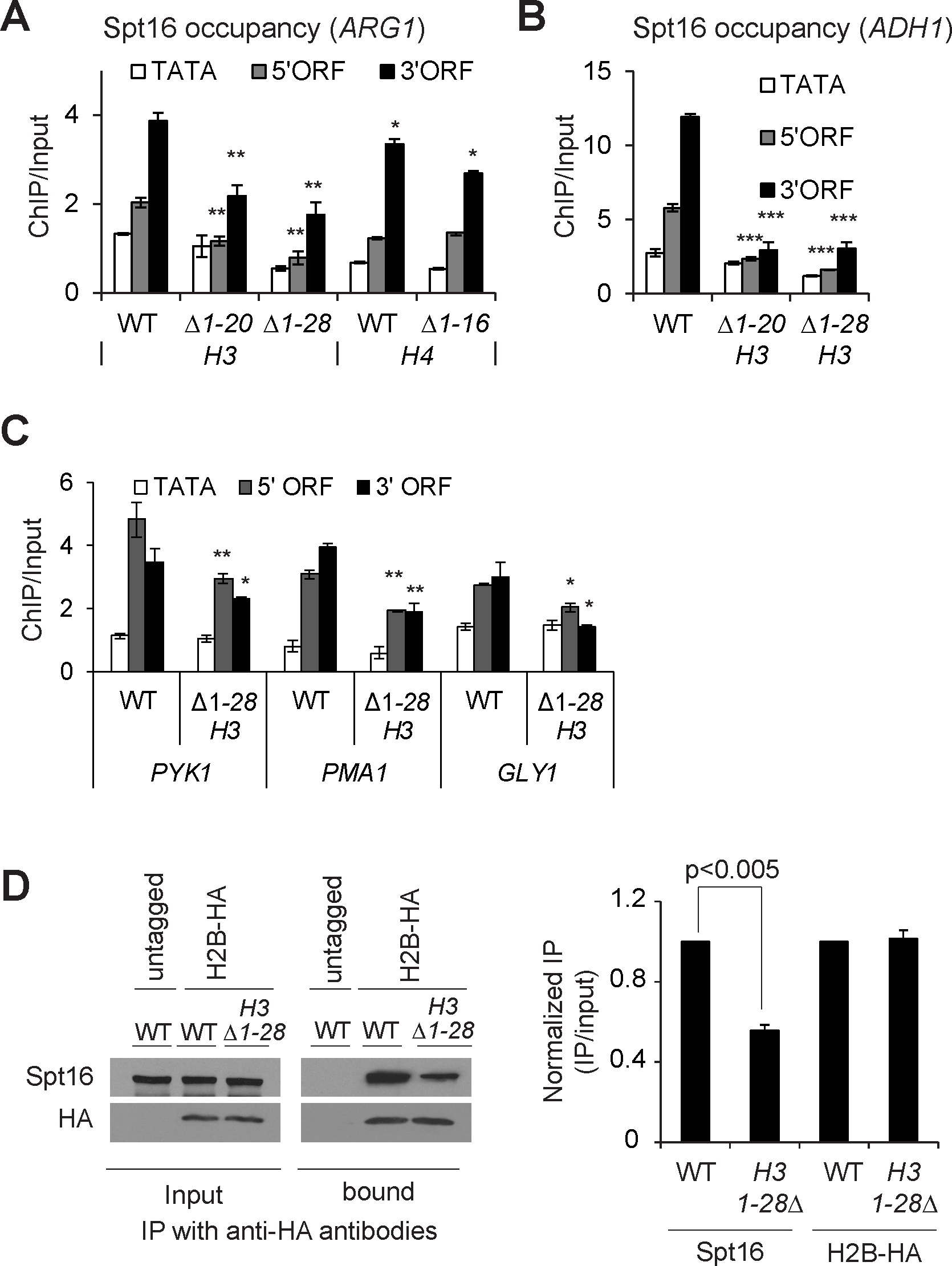
The H3 N-terminal tail promotes FACT recruitment. A-B) ChIP occupancy of Myc-tagged Spt16 in WT and mutants lacking the H3 N-terminal tail residues 1-20 (*H3*Δ*1-20*) or 1-28 (*H3*Δ*1-28*) and 1-16 residues of H4 (*H4*Δ*1-16*) at *ARG1* (A) and in H3 mutants at *ADH1* (B). Graphs show mean and SEM. * represents a p-value < 0.01, ** represents a p-value < 0.001, and *** represents a p-vlaue < 0.0001. C) ChIP occupancies of Myc-tagged Spt16 at the indicated genes in the WT and the H3 tail mutant, *H3*Δ*1-28.* * represents a p-value < 0.01, ** represents a p-value < 0.001. D) Whole-cell extracts prepared from HA-tagged H2B WT and *H3*Δ*1-28* strains were pulled-down with the anti-HA beads, and the immunoprecipitates were subjected to western blot using antibodies against Spt16 and HA. Untagged WT was used as a control. The representative blot is shown on the left, and the quantified data on the right.

To further examine the role of the H3 tails in promoting FACT association with chromatin, we performed coimmunoprecipitation assay. The HA-tagged histone H2B (H2B-HA) efficiently pulled-down Spt16 from the whole cell extracts (WCEs) prepared from HA-tagged WT cells but not from untagged cells (Figure 1D, left). We also observed a reduced Spt16 pull-down from the *H3*Δ*1-28* WCEs (~50 %; Figure 1D, right). No such reduction in Spt16 occupancy was seen in the H4 tail deletion mutant (*H4*Δ*1-16;* Figure S1C), supporting the idea that the H3 tail promotes Spt16 association with chromatin *in vivo.* However, the extent to which the H3 tail contributes in this process may be variable, as observed by the differences of Spt16 occupancies at the different genes in the H3 mutant (Figure 1A-1C).

### Acetylation of the H3 tail promotes FACT occupancy at *ADH1* and *ARG1* genes

After observing that the H3 tail contributes to Spt16, we examined the role for posttranslational modifications on the H3 tail in regulating FACT localization to transcribed genes. The H3 tail lysines, K4 and K36, are methylated by Set1 and Set2, respectively (Briggs *et al.* 2001; Strahl *et al.* 2002).To examine, whether the reduction in Spt16 occupancy in the H3 tail deletion mutant is due to the loss of H3 methylation, we measured Spt16 enrichment in the *set1*Δ/*set2*Δ double mutant. Comparable occupancies of Spt16 were observed in the ORFs of *ARG1* and *ADH1* genes in WT and the *set1*Δ/*set2*Δ mutant (Figure 2A). Furthermore, we found very similar signal for coimmunoprecipitation of Pol II with Spt16 in WT and *set1*Δ/*set2*Δ WCEs (Figure 2B), suggesting that H3 methylation is likely dispensable for maintaining WT level of FACT occupancy, at least, at these two genes. The H3 and H4 tails are also acetylated by Gcn5-containing SAGA and Esa1-containing NuA4 histone acetyltransferase (HAT) complexes, respectively. Accordingly, a *gcn5*Δ*/esa1ts* double mutant elicits strong reductions in H3 and H4 acetylation (Ginsburg *et al.* 2009). The *gcn5*Δ*/esa1ts* mutant, grown at the permissive temperature 25°C (represented as *gcn5*Δ) or at the non-permissive temperature 37°C to inactivate Esa1 (*gcn5*Δ*/esa1ts)*, produced comparable reductions in Spt16 occupancy in the *ARG1* and *ADH1* ORFs (Figure 2C and Figure S2B). In contrast, both WT and the HAT mutant displayed comparable Pol II occupancies at *ARG1* and *ADH1* (Figure 2D, and Figure S2A). To examine whether FACT occupancy correlates to H3 acetylation levels, we determined Spt16 enrichment at *ARG1* in a histone deacetylase mutant, *rpd3*Δ*/hos2*Δ. Interestingly, Spt16 occupancy was not reduced in the histone deacetylase mutant *rpd3*Δ*/hos2*Δ (Figure 2E). This is surprising given that H3 acetylation levels were shown to be elevated in this mutant (Govind *et al.* 2010), and that our current results reveal diminished FACT occupancy in the HAT mutant. It is possible that the WT level of histone acetylation is sufficient for normal FACT occupancy, suggesting that histone acetylation, but not deacetylation, plays a role in FACT recruitment/retention in coding regions. As such, any further increase does not necessarily increase FACT occupancy. Taken together, these results suggest a potential role for the acetylated H3 tail in promoting FACT occupancy in the transcribed coding regions.

**Figure 2:**
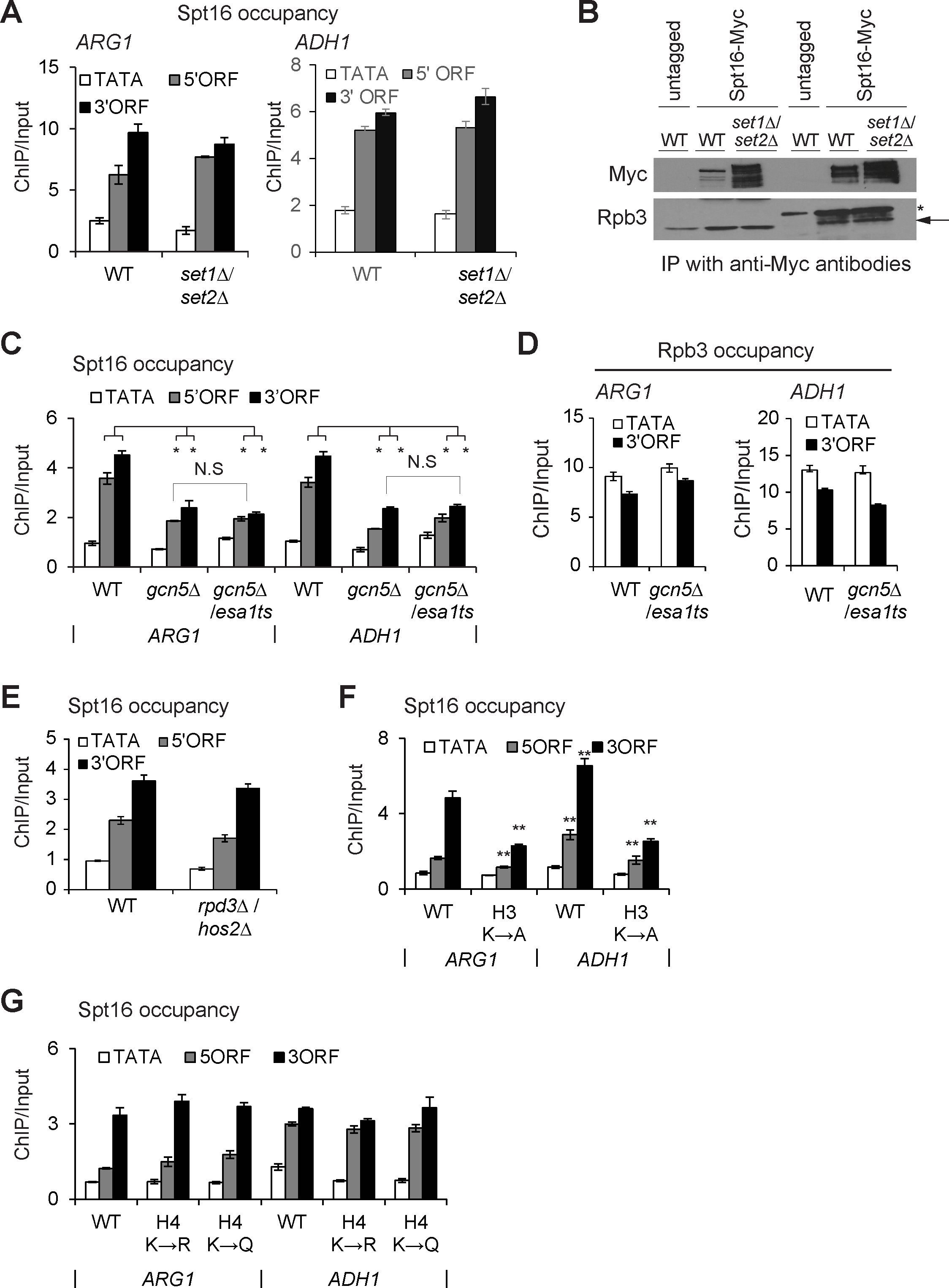
Role of the acetylated H3 tail in modulating Spt16 occupancy. A) ChIP enrichment of Spt16-Myc at *ARG1* in WT and histone methyltransferase mutant *set1Nset2A.* * represents a p-value < 0.001, and ** represents a p-vlaue <0.0001. B) Whole-cell extracts prepared from Spt16-Myc tagged WT and *set1*Δ/*set2*Δ strains were immunoprecipitated with anti-Myc antibodies, and the immunoprecipitates were analyzed by western blot to detect signals for Myc and Rpb3. Untagged WT was used as a control. Rpb3 band in immunoprecipitated samples is shown by an arrow and the IgG heavy chain by an asterisk (*). C) Spt16-Myc ChIP occupancy in WT and *gcn5*Δ*/esa1* histone acetyletransferase (HAT) mutant. Spt16-Myc occupancies were measured by ChIP at *ARG1* and *ADH1* in a *gcn5*Δ*/esa1ts* strain grown either at 25°C (represented as *gcn5Δ*) or at 37°C to inactivate Esa1 (*gcn5*Δ*/esa1*). Graphs show mean and SEM. * represents a p-value < 0.001, and ** represents a p-vlaue <0.0001. Spt16 occupancy differences between *gcn5Δ* and *gcn5*Δ*/esa1ts* were not significnat (N.S). D) Rpb3 occupancies in WT and *gcn5*Δ*/esa1ts* at *ARG1* and *ADH1.* E) Spt16 occupancies in WT and HDAC mutant *hos2*Δ*/rpd3Δ* at *ARG1.* F-G) Spt16-Myc enrichments at *ARG1* and at *ADH1* in histone H3 (F) and H4 tail mutants (G). H3 K4, K9, K14, K18 substituted to alanine, H3K→A; H4 K5, K8, K12 and K16 substituted to arginine; H4K→R, or to glutamine H4K→Q. Graphs show mean and SEM. ** represents a p-vlaue <0.0001.

To provide additional proof for the role of histone acetylation, we examined Spt16 occupancy in the H3 and H4 tail point mutants. The H3 mutant (K4, K9, K14, K18 substituted to alanine; H3K→A) displayed reduced Spt16 occupancy in the ORFs of both *ARG1* and *ADH1* genes (Figure 2F). However, only minimal changes in Spt16 occupancy were observed in the H4 mutant (K5, K8, K12 and K16 substituted to arginine; H4K➔R, or to glutamine H4K➔Q) (Figure 2G; H4K➔A mutant exhibits a lethal phenotype). While Spt16 interacts with both H3 and H4 N-terminal tails, and the histone tail mutants impair FACT function (Biswas *et al.* 2006; VanDemark *et al.* 2008), our results suggest that acetylation of H3 tail makes a greater contribution to the FACT occupancy in the coding regions.

### FACT and Spt6 are required for transcription genome-wide

Microarray analyses examining transcription defects have revealed that the loss of Spt16 function leads to aberrant transcription at many genomic locations, including within coding regions (Cheung *et al.* 2008; van Bakel *et al.* 2013). While it is evident that Spt16 functions to suppress wide-spread cryptic and anti-sense transcription, the role of FACT in regulating Pol II occupancy in coding regions is not well understood at a genome-wide scale. Gene-specific studies have suggested that FACT regulates transcription at the initiation step (Biswas *et al.* 2006; Jimeno-Gonzalez *et al.* 2006; Duina *et al.* 2007). Additionally, FACT, in cooperation with H2B ubiquitination, is important for restoration of chromatin in the wake of Pol II elongation (Fleming *et al.* 2008). As mentioned earlier, similar to the FACT complex, Spt6 (a H3/H4 chaperone) is localized to the coding regions of strongly transcribed genes (Mayer *et al.* 2010; Ivanovska *et al.* 2011; Perales *et al.* 2013; Burugula *et al.* 2014) and is also important for suppressing aberrant transcription (Kaplan *et al.* 2003; Cheung *et al.* 2008; van Bakel *et al.* 2013). To compare the impact of FACT and Spt6 on transcription, we utilized strains in which the expression of *SPT16* or *SPT6* was under the control of a tetracycline repressible promoter (*SPT16-TET* and *SPT6-TET).* These promoters can be repressed by growing cells in the presence of doxycycline (dox).

To rule out unexpected consequences of replacing the endogenous promoter with the TET-promoter, we first compared Spt16 and Spt6 protein levels in TET-strains and BY4741 (S. *cerevisiae* WT strain). Spt16 and Spt6 protein levels in the untreated (no dox; ND) *SPT16-TET* and *SPT6-TET*, respectively, were very similar to those detected in the BY4741 cells (Figure 3A, top panel). As expected, treating *SPT16-TET* and *SPT6-TET* cells with dox led to reduced expression of Spt16 and Spt6, respectively (Figure 3A, bottom panel). We noted that Spt16 was depleted to a greater extent than Spt6 upon dox-treatment. Since Spt16 mutants have been shown to cause cell cycle defects (Prendergast *et al.* 1990), we also measured the level of budded and unbudded cells in BY4741, *SPT16-TET* and *SPT6-TET* dox-treated cells. We did not find any significant increase in the number of budded and unbudded cells under Spt6 or Spt16 depleted conditions (Figure S3A), suggesting that depleting Spt16 or Spt6, under the experimental conditions employed, elicit minimal cell cycle defects. Moreover, the TET-strains grown in the presence or absence of dox, exhibited similar viability (Figure 3B). Altogether, these results indicate that the untreated *SPT16-TET* and *SPT6-TET* cells behave similar to BY4741.

**Figure 3:**
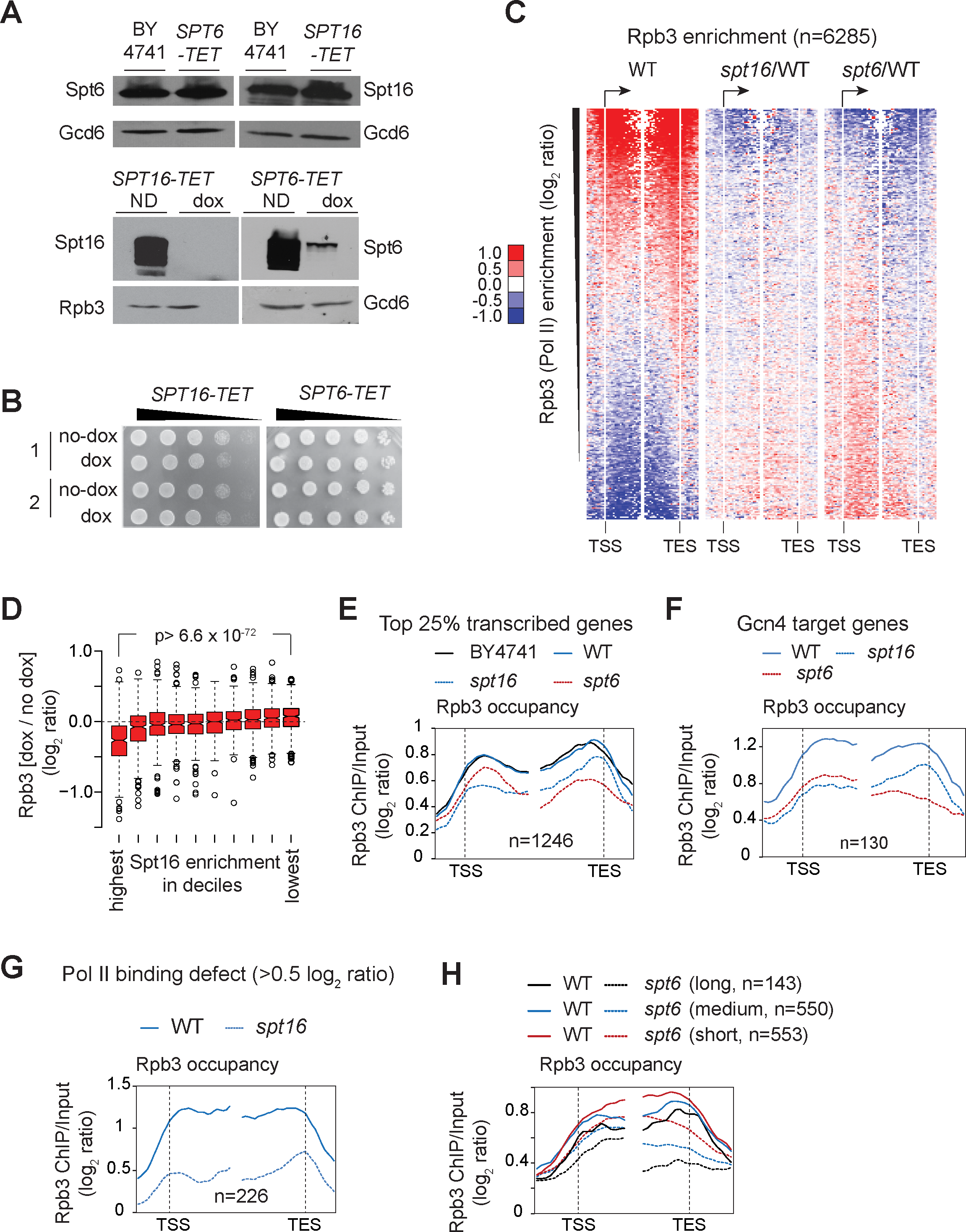
Effect of depleting Spt16 and Spt6 on Pol II occupancy genome-wide. A) *BY4741, SPT16-TET* and *SPT6-TET* cells were grown in SC media and induced by SM. Western blots show Spt16 and Spt6 protein levels in BY4741, *SPT16-TET* and *SPT6-TET* strains without dox-treatment (top panel). The Spt16 and Spt6 levels in untreated (ND) and dox-treated TET-strains are shown (bottom panel). B) *SPT16-TET* and *SPT6-TET* cells grown and subcultured in SC media, with and without doxycycline (dox), were collected, serially 10-fold diluted, and spotted on SC plates. Growth for the two cultures of each *SPT16-TET* (left) and *SPT6-TET* (right) with and without dox (no-dox) treatment are shown. C) Heat-maps depicting genome-wide Rpb3 (Pol II) enrichment in *SPT16-TET* (without dox; WT) (left), and changes in Rpb3 occupancies in dox-treated *SPT16-TET* (spt16/WT) (middle) and *SPT6-TET* (*spt6*/WT) (right). Genes were sorted from highest to lowest ORF Rpb3 enrichment in WT cells. D) Box-plot showing the changes in Rpb3 occupancy according to the Spt16 enrichment in deciles. The average ORF occupancy of Rpb3 in *SPT16-TET* dox-treated and untreated cells was determined, and the change in Rpb3 occupancy on depleting Spt16 (dox/no dox) was calculated for each gene. Genes were then grouped into deciles according to the Spt16 occupancy in WT cells. The first decile shows the highest Spt16 occupancy and the 10^th^ shows the lowest. E) Rpb3 occupancy profiles for the top 25% Pol II-occupied genes (n=1246) in untreated *BY4741* and *SPT16-TET* (WT), and in dox-treated *SPT16-TET (spt16*) and *SPT6-TET (spt6*) cells are shown. F) Rpb3 occupancy profiles in untreated *SPT16-TET* (WT), and dox-treated *SPT16-TET (spt16*) and *SPT6-TET (spt6*) cells at the Gcn4 targets genes enriched in the top 25% Pol II-occupied genes. G) Rpb3 occupancy profile at the genes eliciting reduction in Rpb3 occupancy ≥ 0.5 log_2_ ratio (ChIP/input) on depleting Spt16 in WT and *spt16.* H) The top 25% Pol II-occupied genes were grouped on the basis of their gene-length and Rpb3 occupancy profiles for the long (>2 kb), medium (1-2 kb), and short (0.5-1 kb) are shown in the WT and Spt6-depleted cells.

To examine the effects of depleting Spt16 and Spt6 on transcription, we determined Rpb3 occupancy in untreated cells (*SPT16-TET;* referred to as WT hereafter), and dox-treated *SPT16-TET (spt16*) and *SPT6-TET (spt6*) cells by ChIP-chip. We observed a strong correlation (Pearson correlation, r= 0.93) between the Pol II occupancies in the dox-untreated *SPT16-TET* and BY4741 strains, genome-wide, which further indicated that replacing the endogenous *SPT16* promoter with the TET-promoter does not adversely affect transcription. We also determined Spt16 occupancy genome-wide by ChIP-chip and found that Spt16 occupancy in coding regions strongly correlated with Pol II occupancy (Pearson correlation, r=0.85), in agreement with previous studies (Mayer *et al.* 2010).

The heat-maps depicting changes in Rpb3 enrichment (*spt*16/WT) showed diminished ratios in coding regions of the genes displaying the greatest Rpb3 enrichments in WT cells (Figure 3C). Consistent with a strong correlation between Spt16 and Rpb3 occupancies, the genes with the highest Spt16 enrichments showed greatest Rpb3 reductions (Figure 3D).

We also noted that deletion of Spt16 (and Spt6, described later) also revealed an increase in Pol II occupancies at those genes, which otherwise show very poor enrichment ratios in WT cells. Given that our ChIP-chip normalization was performed without spike-in control, this apparent increase in Rpb3 occupancy is unlikely to be biologically relevant.

To further analyze the impact of depleting Spt16 on Pol II occupancy, we selected the top 25% genes showing greatest Rpb3 occupancy in WT cells (n=1246). Nearly identical profiles for Rpb3 occupancy were observed in WT and BY4741 at the metagene comprised of these transcribed genes (Figure 3E). However, Spt16 depletion evoked reduction in Pol II occupancy, primarily in coding region of these sets of genes. Although reduced Pol II occupancy was observed throughout the coding region, Pol II occupancy defect was more pronounced in the 5’ end. Intriguingly, Pol II occupancy appeared to increase towards the 3’ end, relative to the 5’ end, in the Spt16-depleted cells. Considering that if Pol II fails to clear the promoter or disengages near the 5’ end under limited Spt16 availability, then Pol II occupancy should have been low uniformly across coding region. The relative increase in Pol II occupancy towards the 3’ end may reflect increased cryptic transcription events that accumulate Pol II from the 5’ to 3’ end. Such an explanation would be consistent with the established role of histone chaperones in suppressing cryptic transcription (Kaplan *et al.* 2003; Cheung *et al.* 2008; van Bakel *et al.* 2013). A previous study utilizing microarray predicted 960 and 1130 genes to have cryptic transcription in *spt6-1004* and *spt16-197* mutants (Cheung *et al.* 2008). We found that only 154 genes, predicted to express cryptic transcripts in the previous study, were among the 1246 genes exhibiting high-levels of Pol II occupancy (Figure S3B). Replotting Pol II occupancy data after excluding these 154 genes (Figure S3C) displayed profiles similar to that observed in Figure 3E. This analysis suggests that it is unlikely that differences in Pol II profiles observed under Spt6 and Spt16 depleted condition is a result of the presence of genes expected to display cryptic transcription. It is possible that Pol II pausing and queuing Pol II in the 3’ end superimposed on elongation defects at the very 5’ end could result in increased Pol II occupancy towards the 3’ end on depleting Spt16.

Given that Gcn4 target genes are activated under the growth conditions used (see Materials and Methods), we additionally analyzed the effect of depleting Spt16 on transcription of Gcn4 targets. 130 Gcn4-regulated genes were enriched among the top 1246 transcribed genes. These genes also displayed a greater reduction in Pol II occupancy at the 5’ end in Spt16-depleted cells (Figure 3F). A similar profile for Rpb3 occupancy defect was observed in ribosomal protein genes, which are among the highly transcribed genes (*data not shown*). We identified 226 genes showing reduction in Pol II occupancy ≥ 0.5 log_2_ ratio (ChIP/input) in Spt16-depleted cells (Figure 3G). Interestingly, these genes were enriched among the top 10% transcribed genes (p-value=10^−117^), suggesting that depletion of Spt16 imparts a significant effect on Pol II occupancy at highly expressed genes. Collectively, these results show that Pol II occupancy in coding region is differentially affected by the loss of Spt16 and implicates FACT in regulating global transcription.

Next, we analyzed Pol II occupancy in the Spt6 depleted cells. Interestingly, at the top 25% of Pol II-occupied genes, Spt6-depletion elicited a greater reduction in Rpb3 occupancy towards the 3’ end (Figures 3C (right) and 3E), consistent with previous studies showing the greatest Spt6 enrichments in the 3’ ends of transcribed genes (Perales *et al.* 2013; Burugula *et al.* 2014). A 5’ to 3’ bias in Pol II occupancy was also evident at the Gcn4-targets (Figure 3F) and at 238 genes, which showed a reduction in Pol II occupancy ≥ 0.5 log_2_ ratio (ChIP/input) (Figure S3D). A progressive reduction in Pol II occupancy in the 5’ to 3’ direction, in Spt6-depleted cells, suggests that Spt6 may regulate Pol II processivity. Alternately, given the role of Spt6 activity in the 3’—mRNA processing, diminished Spt6 levels could also result in reduced Pol II occupancy at the 3’ end (Kaplan *et al.* 2005). To distinguish between these two possibilities, we analyzed Rpb3 occupancy at the top 25% transcribing genes based on their gene length. All three groups of genes, long (> 2 kb), medium (1-2 kb) and short (0.5-1 kb), showed a 5’ to 3’ bias in Pol II occupancies (Figure 3H). Interestingly, however, long and medium genes displayed a greater reduction in the 3’ end compared to the short genes (0.5-1 kb) (Figure 3H and S3E). A simpler explanation for this observation is that Spt6 promotes Pol II elongation, in agreement with previous studies (Endoh *et al.* 2004; Ardehali *et al.* 2009; Perales *et al.* 2013). A 3—mRNA processing defect (Kaplan *et al.* 2005) would be expected to produce similar reductions in Pol II occupancies at the 3’ end irrespective of gene length. Collectively, our data suggest that loss of FACT and Spt6 functions produces distinct effects on Pol II occupancy. However, the role of FACT and Spt6 in suppressing aberrant transcription, to some extent, could have an effect on Pol II occupancy under depletion conditions.

### FACT and Spt6 differentially impacts transcription and histone occupancy

To further address the functional overlap between FACT and Spt6, we examined genes, which showed a reduction in Pol II occupancy (≤ −0.5 log_2_ ratio) upon depleting these factors. We found a significant overlap between the genes exhibiting Pol II fold-change log_2_ ≥ 0.5 upon depleting either Spt16 or Spt6 (p-value = 5.1 × 10^−76^, n= 111) (Figure 4A). The genes showing Pol II occupancy defects upon depleting either Spt16 or Spt6 (common; n=111) exhibited, on average, higher Pol II occupancy (in WT cells) than those genes which showed defects only after depleting either Spt16 or Spt6 (unique) (Figure 4B). This observation suggests that strongly transcribed genes may need full functions of Spt6 and Spt16 for a high-level of transcription. It is also interesting to note that while Spt6 depletion was less efficient compared to that of Spt16, it nonetheless evoked very similar Pol II occupancy defects on these genes (Figure 4C). In contrast, Rpb3 profiles at the genes uniquely affected by Spt6 or Spt16 depletion showed a distinct cohort behavior. Greater reduction in Rpb3 occupancy was observed across the ORF of the Spt16-unique genes (n=115) upon depletion of Spt16 than of Spt6 (Figure 4D). Unlike the ‘common genes’ these genes tolerated a moderate loss of Spt6. Likewise, Spt6-unique genes (n=217) displayed a stronger Pol II occupancy defect after depleting Spt6 than Spt16 (Figure 4E), which was depleted to a larger extent. Considering that Spt6 exhibits higher occupancy towards the 3’ ends (Mayer *et al.* 2010; Perales *et al.* 2013; Burugula *et al.* 2014), we analyzed distribution of the gene-length in the three classes of genes. The longer genes were enriched in the Spt6-unique and common genes, whereas Spt16-unique genes were shorter (Figure S4A and S4B). Thus, it appears that the differential effect of Spt16 and Spt6 depletion could partly be due to differences in their localization patterns over the coding sequences.

**Figure 4:**
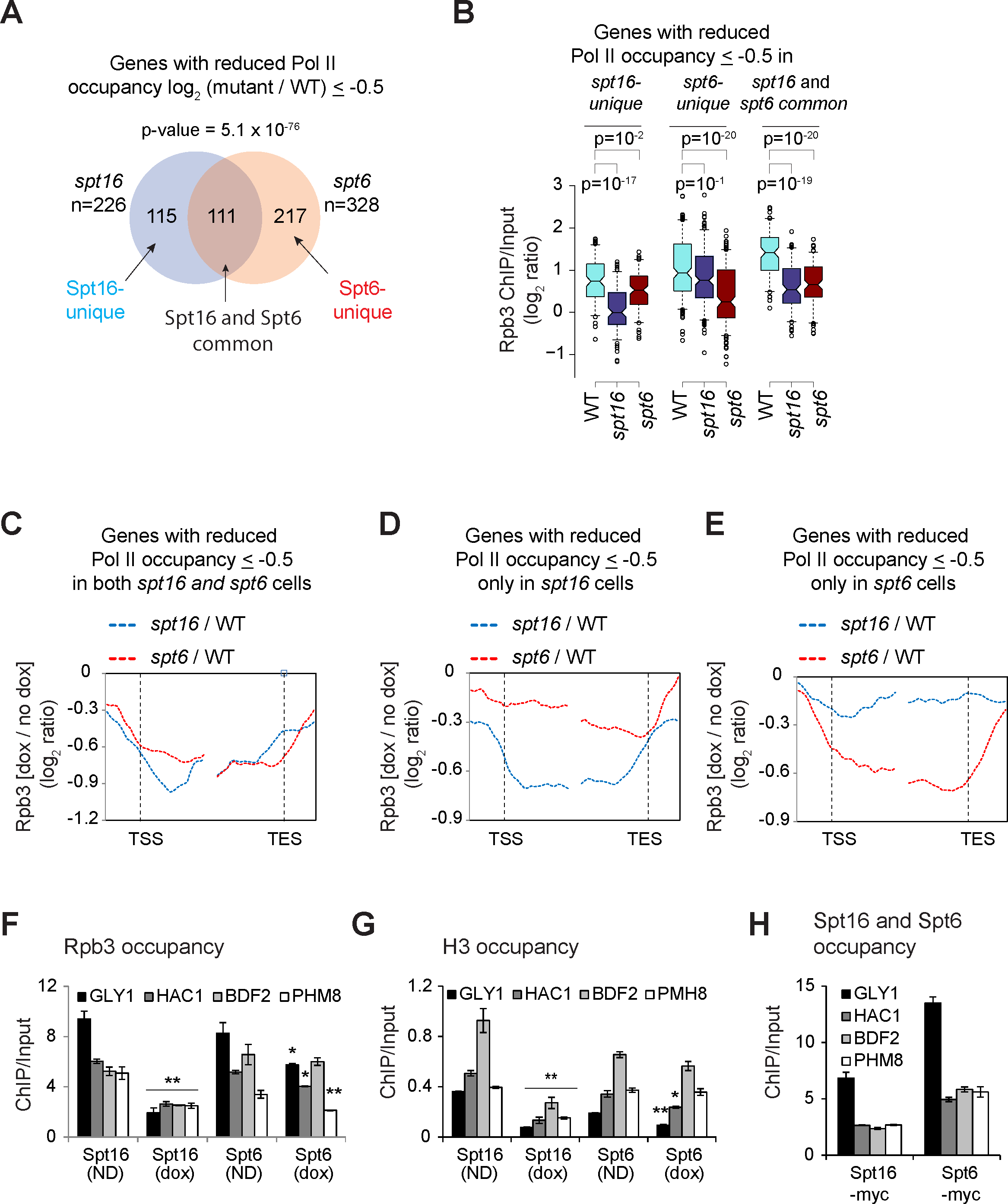
FACT promotes nucleosome reassembly. A) Venn diagram showing overlap among the genes which displayed Rpb3 occupancy defect > 0.5 log_2_ ratio (ChIP/Input) in Spt16 and Spt6 depleted cells. p-value for the overlap is shown. B) Box-plot showing Rpb3 average ChIP-chip enrichments at Spt16-unique, Spt6-unique, and Spt16 and Spt6 common genes in WT, Spt16-depleted and Spt6-depleted cells. C-E) Metagene analysis for changes in Pol II enrichment under Spt16 and Spt6 depeleted cells, for genes showing reduction > 0.5 log_2_ ratio (ChIP/Input) under Spt16 or Spt6 depleted condition (C; common), only under Spt16 depletion (D) only under Spt6 depletion (E). F-H) ChIP occupancies of Rpb3 (F), histone H3 (G), and of Spt6 and Spt16 in 5’ ORFs of indicated genes in *SPT16-TET* and *SPT6-TET* untreated (ND) or dox-treated (dox) cells. * represents a p-value < 0.05, and ** represents a p-vlaue <0.001.

We further examined the differential effect of depletion by determining Rpb3 ChIP occupancy at four genes, which showed comparable Spt6, and Spt16 occupancy in our ChIP-chip experiments. Pol II occupancies in the ORFs of *GLY1, HAC1, BDF2*, and *PHM8* were substantially reduced (~2-5 folds) upon Spt16 depletion (Figure 4F). By contrast, only a moderate to negligible reduction was observed after depleting Spt6. For example, Rpb3 was reduced by less than 1.5 folds at *GLY1, HAC1, PMH8* upon Spt6 depletion. Similarly, we found that histone H3 occupancy was more severely reduced in 5’ ORFs of these genes upon depleting Spt16 than upon Spt6 (Figure 4G). Reduced H3 occupancies upon depletion of Spt16 and Spt6 are consistent with their role in histone reassembly. Bigger reductions in Pol II and H3 occupancies were not due to higher Spt16 occupancies at these genes compared to that of Spt6 (Figure 4H). Collectively, these results suggest that Spt6 and Spt16 may help promote transcription in a gene-specific manner.

### Spt6 promotes histone eviction to promote transcription

To further investigate whether Spt6 and FACT cooperate in promoting transcription, we deleted the Spt6 tandem SH2 domain (tSH2; 202 residues from C-terminus), which mediates Spt6 recruitment, genome-wide (Mayer *et al.* 2010; Burugula *et al.* 2014), in the *SPT16-TET* background (*SPT16/spt6*Δ*202).* As expected, treating the *SPT16/spt6*Δ*202* mutant with dox resulted in reduced Spt16 protein levels (Figure S5A). We then determined Rpb3 and H3 occupancies by ChIP-chip in untreated (*SPT16/spt6*Δ*202*) and dox-treated (*spt16/spt6*Δ*202*) cells, and compared the occupancies with that of the untreated *SPT16-TET* cells (WT). The changes in Rpb3 occupancy in untreated *SPT16/spt6*Δ*202* were significantly anti-correlated with the Rpb3 occupancy in the WT cell (r= −0.81) (Figure 5A), indicating a strong requirement of Spt6 in transcription, genome-wide. A modest but statistically significant reduction in Rpb3 occupancy was observed in dox-treated, *spt16/spt6*Δ*202*, cells compared to the untreated cells (for the first decile, p =5.2 × 10^−59^), indicating that these factors cooperate to promote efficient transcription (Figure 5B).

**Figure 5:**
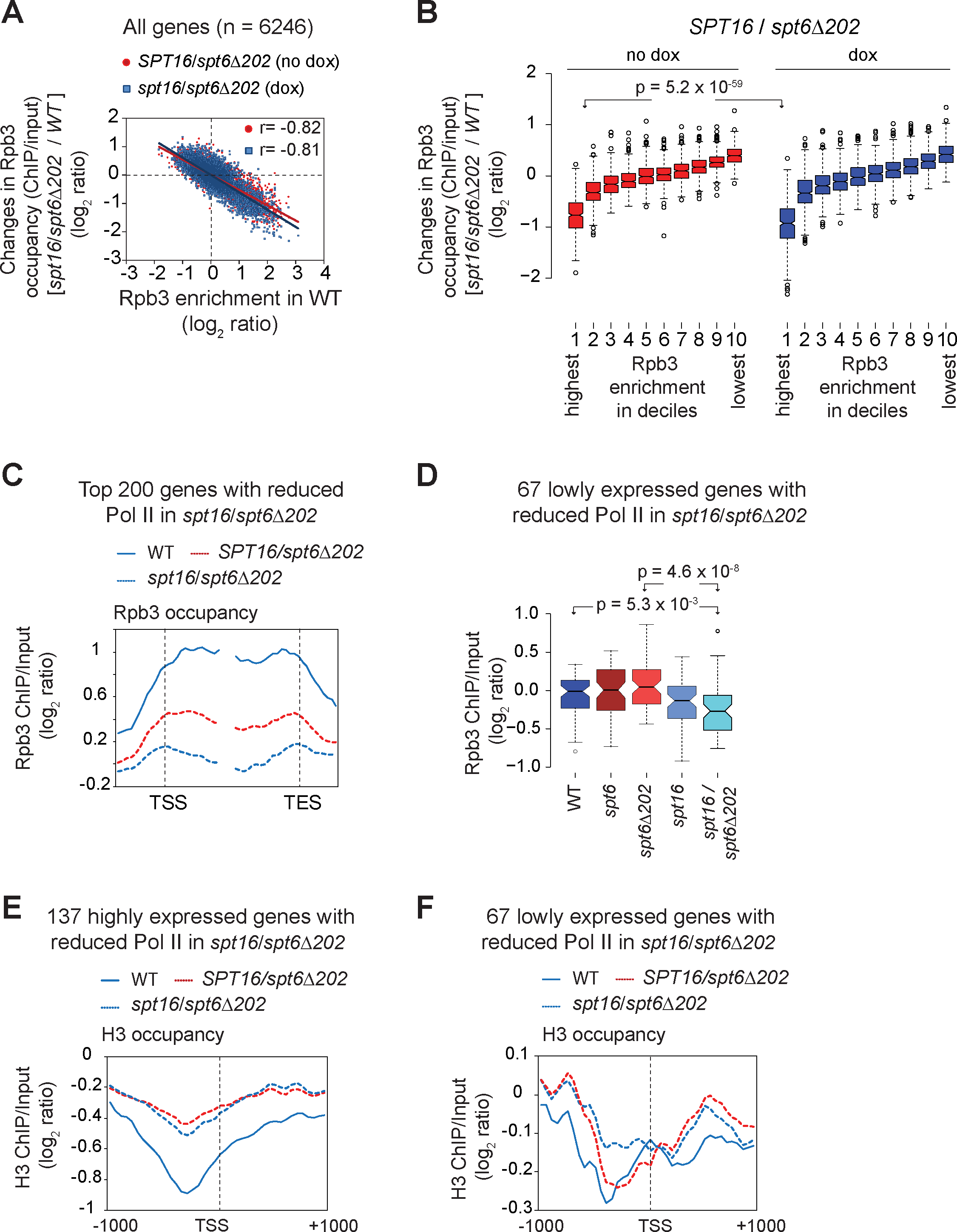
Spt6 promotes histone eviction and transcription. A) The changes in Rpb3 occupancies (log_2_ ratio, ChIP/input) in the untreated (no dox) and treated (dox) *spt16/spt6*Δ*202* cells relative to occupancies in the untreated *SPT16-TET* (WT) are plotted against the Rpb3 ORF occupancies in WT. Pearson correlations are shown. B) Box-plot showing changes in Rpb3 occupancy in *SPT16/spt6*Δ*202* (dox and no dox) relative to the *SPT16-TET* (no dox) WT cells, at the genes grouped in deciles on the basis of the average ORF Rpb3 occupancies observed in WT cells. The first decile shows the highest Rpb3 occupancy and the 10^th^ shows the lowest. C) Top 200 genes showing the greatest reduction in Pol II occupancy in dox-treated *SPT16/spt6*Δ*202* relative to the untreated cells were selected based on the averge Rpb3 enrichments in coding regions. Pol II occupancy profile for these genes is plotted for WT, *SPT16/spt6*Δ*202* and *spt16/spt6*Δ*202.* D) Box-plot showing average Rpb3 enrichment (log_2_ ratio, ChIP/input) for 67 genes of the 200 genes. These genes were not among the top 25% genes showing greatest Rpb3 enrichment in WT cells. p-values are shown. E-F) H3 occupancy profile for 137 highly expressed genes (E) and at 67 lowly expressed genes among the 200 genes showing greatest reduction in *Spt16/spt6*Δ*202* dox-treated compared to untreated cells (F).

Rpb3 profiles in *spt16/spt6*Δ*202* mutant revealed greatly diminished occupancy across the coding regions of the top 25% Pol II-occupied genes (Figure S5B). We noted that untreated *SPT16/spt6*Δ*202* elicited greater reduction in Pol II occupancy than observed upon depleting Spt6 (compare Figures S5B and 3E). This could be a result of an incomplete depletion of Spt6, and thus, enough Spt6 remains after dox-treatment to promote transcription, albeit at reduced efficiency. It is also possible that the tSH2 domain has additional roles in modulating Spt6 function in a manner that deleting tSH2 evoked a stronger Pol II occupancy defect than depleting Spt6. Nonetheless, these results indicate the importance of Spt6 in stimulating high-level transcription, genome-wide. However, depleting Spt16 in *SPT16/spt6*Δ*202* cells produced only modest reductions in Pol II occupancy at the top 25% Pol II occupied (Figure S5B). This observation raises a possibility that FACT may require certain aspects of Spt6 function to stimulate transcription. Such an explanation is consistent with the observation that Spt16 depletion elicited reduced Pol II occupancy in otherwise WT strain (Figures 3D-G).

To further address functional cooperation between Spt16 and Spt6, we focused on the genes eliciting greater reduction in dox-treated *spt16/spt6*Δ*202* than in untreated cells. The top 200 genes showing the greatest Pol II occupancy defect revealed that depleting Spt16 in *spt6*Δ*202* background significantly reduced Rpb3 occupancy across the coding region (Figure 5C). 137 of these 200 genes were among the top 25% expressed genes (n=1246) in WT cells. While these 137 genes displayed greater Pol II occupancy defects in *SPT16/spt6*Δ*202* than observed in Spt6 or Spt16 depleted cells, the greatest defect was observed in the double mutant (Figure S5C). These observations suggest that FACT and Spt6 may act synergistically to stimulate transcription of a subset of highly expressed genes.

We also examined the 63 lowly expressed genes among the 200 genes, which showed greater reduction in the double mutant. While no significant reduction in Pol II occupancy was seen either in *SPT16/spt6*Δ*202* mutant or in Spt6 and Spt16 depleted cells, occupancy was significantly reduced in the double mutant (Figure 5D). This suggests that FACT may redundantly act with Spt6 in promoting transcription of a subset of lowly expressed genes. More sensitive methods may be needed to fully comprehend the extent to which Spt6 and FACT coordinate transcription of lowly expressed genes, especially considering the technical challenges in accurately measuring Pol II occupancies at genes expressed at very low levels.

We next examined histone occupancy changes in *spt16/spt6*Δ*202.* We found that the 137 highly expressed genes displayed increased histone occupancy across the metagene (Figure 5E). The greatest increase in histone occupancy was observed slightly upstream of the TSS. Similar profile was observed for the top 25% genes (n= 1246; data not shown). These results suggest that in addition to reassembling histones, Spt6 may play a role in stimulating removal of histones during transcription. Since histones pose a significant barrier to transcription (Teves *et al.* 2014), elevated histone occupancy could explain a robust transcription defect observed at transcribed genes in *SPT16/spt6*Δ*202* cells. However, despite a greater reduction in Pol II occupancy in dox-treated cells, no significant increase in H3 occupancy was observed in the double mutant compared to the single. One possible explanation for these results is that FACT restructures chromatin to enhance transcription in a manner independent of histone disassembly, as proposed earlier (Formosa 2012). In contrast, the histone occupancy profile for the 67 genes (low transcribed) was distinct from the highly expressed genes in that elevated histone occupancy near the TSS was observed only in the *spt16/spt6*Δ*202* double mutant, but not in the single (*SPT16/spt6*Δ*202*) (Figure 5F). However, in both treated and untreated cells, elevated histone occupancies were observed downstream of the TSS. This observation is in agreement with the Pol II occupancy defects, which were also observed only in the double mutant (Figure 5F). Increased histone occupancies near the TSS and in coding regions are consistent with the idea that histones pose a significant barrier to transcription and suggest a role for FACT and Spt6 in modulating histone occupancy.

## DISCUSSION

In this study, we have examined the role of two highly conserved histone chaperones, Spt16 and Spt6, in regulating genome-wide transcription and histone occupancy, under amino acid starvation conditions. Spt6 recruitment to coding regions is stimulated by the phosphorylated Pol II CTD and by HDACs Rpd3 and Hos2 (Mayer *et al.* 2012; Burugula *et al.* 2014). However, the mechanism by which the FACT complex associates with chromatin is not well understood. Our results showing diminished occupancy of Spt16 at the ORFs of *ARG1* and *ADH1* and reduced interaction with nucleosomes in mutants lacking the H3 tail (Figure 1A-1C), suggests a role for the H3 tail in promoting FACT association with transcribed regions. Our results additionally suggest that histone acetylation enhances the ability of the FACT complex to associate with transcribed ORFs, since deleting Gcn5 (a H3 HAT) or mutating the H3 tail lysine residues significantly dampened Spt16 enrichment in coding sequences of *ARG1* and *ADH1* (Figures 2C and 2F). These results provide *in vivo* evidence for the previous studies showing impaired FACT binding to the nucleosomes/histones lacking N-terminal tails (VanDemark *et al.* 2008; Winkler *et al.* 2011). Currently, however, which regions of Spt16 are required for histone tail interactions is unclear. While Spt16 N-terminal domain of *S. pombe* was shown to be important for interacting with H3 and H4 tails, in *S. cerevisiae*, this domain was found to be dispensable (VanDemark *et al.* 2008).

Considering that Spt16 exhibits similar affinity towards the acetylated and unacetylated histone peptides *in vitro* (Stuwe *et al.* 2008), it is plausible that acetylation-mediated changes to the nucleosome structure allow FACT to stably bind chromatin at transcribing loci. Consistent with this idea, Gcn5 promotes histone eviction, Pol II elongation, and stimulates recruitment of bromodomain-containing chromatin remodelers, RSC and SWI/SNF (Govind *et al.* 2007; Dutta *et al.* 2014; Spain *et al.* 2014). Additional contacts with core domains of H3/H4 and with H2A/H2B through its C-terminal domain (Winkler *et al.* 2011; Kemble *et al.* 2015) could further stabilize FACT-chromatin interactions, thereby maintaining chromatin in an accessible conformation to promote transcription, and concomitantly aiding in reassembly of evicted histones in the wake of transcription (Jamai *et al.* 2009). Additional factors may act cooperatively with other factors to enhance FACT enrichment. For instance, FACT interacts with the Paf1 complex, and chromatin remodeler Chd1, both of which are enriched in transcribed regions (Krogan *et al.* 2002; Squazzo *et al.* 2002; Simic *et al.* 2003).

Enrichment of FACT in coding regions (Mayer *et al.* 2010) (*data not shown)*, and its ability to promote Pol II transcription through the nucleosomal templates, *in vitro*, (Belotserkovskaya *et al.* 2003; Hsieh *et al.* 2013), strongly suggests a role for FACT in the elongation step of transcription. FACT is linked to the reestablishment of the disrupted chromatin structure in the wake of transcription (Jamai *et al.* 2009), and consequently, in suppressing aberrant transcription by preventing utilization of cryptic promoters (Kaplan *et al.* 2003; Cheung *et al.* 2008; van Bakel *et al.* 2013). Our data indicate that FACT is globally required for promoting transcription in the coding regions. Spt16 deficiency reduced Pol II occupancy in coding regions of highly transcribed genes, including of the ribosomal protein genes and Gcn4 targets (Figure 3). It was previously reported that histone mutants which perturb association of Spt16 (shift in occupancy towards the 3’ end) also reduced Pol II occupancy at the 5’ end of *PMA1* and *FBA1* genes (Nguyen *et al.* 2013). Thus, it seems that impairing FACT activity, through depletion of Spt16 or by altering FACT association in the coding region (Nguyen *et al.* 2013), elicit greater defects in transcription in the early transcribed regions. These results raise a possibility that FACT may help polymerases in negotiating the nucleosomal barrier downstream of the TSS. Such an idea is consistent with biochemical studies showing that FACT relieves Pol II pauses well within the nucleosomes and acts primarily to overcome tetramer-DNA contacts (Bondarenko *et al.* 2006). Considering that the rate of elongation in the early transcribed regions is slower than that observed in the mid or 3’ end regions (Danko *et al.* 2013), our findings suggest that FACT may help slow-moving polymerases to overcome the nucleosomal barrier at the 5’ ends of the transcribed genes. Although, nucleosomes are expected to pose a similar block to elongating Pol II irrespective of their position along coding sequences, nucleosomal impediment to transcription at the distal end may be additionally relieved by factors, such as chromatin remodelers RSC and SWI/SNF, which are enriched in the ORFs of many genes (Dutta *et al.* 2014; Spain *et al.* 2014). Moreover, post-translational histone modifications, which are not uniform across coding regions, may have a differential effect on the FACT activity to stimulate transcription. In support of such a possibility, it was shown that the FACT complex cooperates with H2B ubiquitination to regulate transcription both *in vitro* and *in vivo* (Pavri *et al.* 2006; Fleming *et al.* 2008).

In contrast to FACT, Spt6-depletion diminished Pol II occupancy at the 3’ end (Figures 3E-F and 3H). Lower Pol II occupancy towards the 3’ end could result from a processivity defect and/or from a specific defect at the 3’ end. Our analyses, based on gene-length, favor the idea that Spt6 enhances Pol II processivity, possibly through stimulating nucleosome eviction, or by preventing premature dissociation during Pol II traversal through coding regions. Such an interpretation is consistent with previous studies showing reduced Pol II occupancy at the 3’ end of exceptionally long-genes (Perales *et al.* 2013). Spt6 has been also shown to enhance the rate of Pol II elongation at heat-shock genes in *Drosophila* (Ardehali *et al.* 2009). Therefore, reduced Pol II occupancy near the 3’ ends of yeast genes suggests that Spt6 may function to help elongating polymerase traverse coding regions through multiple mechanisms.

Pol II occupancy in the Spt6 mutant lacking the C-terminal tandem SH2 domain was strongly correlated with Pol II occupancy in the WT cells, indicating a strong requirement for Spt6 in stimulating transcription genome-wide. Even though depleting Spt16 in otherwise WT cells evoked reduction in Pol II occupancy from highly transcribed genes, this surprisingly had only a minor impact on Pol II occupancy at the majority of transcribed genes in *spt6*Δ*202* cells. This suggests that both factors might have overlapping role and that FACT may rely on the certain aspects of Spt6 functions in promoting transcription. It makes sense considering that FACT, and Spt6 are recruited to transcribed regions (Mayer *et al.* 2010; Burugula *et al.* 2014), and that both factors can interact with H2A-H2B and H3-H4 (Mccullough *et al.* 2015). Despite interacting with nucleosomes as well as with histones, unlike Spt6, FACT can reorganize the nucleosome structure in a manner that increases DNA accessibility. This distinct ability of FACT could explain reduction in Pol II occupancy observed in a subset of transcribed genes in *spt16/spt6*Δ*202* double mutant, but not in the single mutant (Figure 5C).

The severe reduction in Pol II occupancy in *spt6*Δ*202* was associated with elevated histone occupancy in both promoter and coding regions, implicating Spt6 in promoting histone eviction to allow efficient transcription. Increased occupancy in the promoter regions observed in the mutant, is in agreement with earlier studies showing that histones are evicted from promoters to allow high-levels of transcription activity (Reinke *et al.* 2001; Dion *et al.* 2007). Loss of Spt6 function in the ΔSH2 mutant could also be responsible for increased histone occupancies, in the coding regions. This idea is supported by studies showing a role for Spt6 in enhancing elongation rate (Ardehali *et al.* 2009) and promoting Pol II processivity ((Perales *et al.* 2013) and Figure 3). However, considering that the Spt6 missing the tSH2 domain can still associate with the ORFs albeit at a reduced level (Mayer *et al.* 2010; Burugula *et al.* 2014), it is plausible that the ORF-associated mutant Spt6 acts in a non-specific manner and impairs transcription leading to elevated histone occupancies across the coding regions. It is also possible that the tSH2 domain modulates Spt6 function such that Spt6 without this domain is impaired for its histone eviction function, leading to strong transcription defects as observed in the *spt6*Δ*202* mutant. The greater Pol II occupancy defect observed upon Spt16 depletion in the tSH2 mutant at a subset of highly transcribed genes (137 genes, Figure S2C) could be explained by the property of FACT to reorganize/destabilize histone-DNA interactions (Xin *et al.* 2009; Formosa 2012) that allows Pol II to negotiate nucleosomal block without requiring eviction of histones.

## ACKNOWLEDGEMENTS

We thank Tim Formosa for providing antibodies against Spt6 and Spt16. We also thank Alan Hinnebusch, Randy Morse and Jeena Kinney for useful discussions and providing valuable comments on the manuscript. CKG is supported by grants from the National Institutes of Health (GM095514), and Center for Biomedical Research (CBR, Oakland University). We also acknowledge the genomic core facility at Michigan State University.

## Legends to the supplemental figures

**Figure S1: Role of the H3 tail in FACT recruitment.**

A) Spt16 ChIP occupancy at *ADH1* in a H4 tail mutant lacking the first 16 residues (*H4*Δ*1-16).*

B) Spt16 and Rpb3 ChIP occupancy in the mutant was normalized against the WT occupancy at each gene and the normalized data is presented for the indicated genes.

C) Anti-HA beads were used to pull down HA-tagged H2B from the whole cell extracts prepared from the untagged WT, the HA-tagged WT and *H4*Δ*1-16* cells. The immunoprecipitates were subjected to western blot using anti-HA and anti-Spt16 antibodies. The representative blot is shown on the left, and the quantified data on the right.

**Figure S2: Role of histone acetylation in recruiting Spt16.**

ChIP occupancies of Rpb3 (left) and Spt16 (right) at *ARG1* in WT, *H3K➔A*, and *gcn5*Δ mutants were determined by real-time (RT) PCR. ChIP occupancy in the *ARG1* 3’ ORF in WT were normalized to *POL1* signals from the same ChIP sample, and normalized again with the *ARG1/POL1* signals for the related input samples. Average data with SEM for three independent experiments is shown.

**Figure S3: Rpb3 occupancy in WT, and Spt16 and Spt6 depleted cells.**

A) A bar-graph showing the bud counts in *SPT16-TET* and *SPT6-TET* cells in the presence or in the absence of doxycycline (dox). The cells were grown exactly as they were for the ChIP-chip experiments. The cells were briefly sonicated before counting budded and unbudded cells under the microscope. The error bars represent standard error of the mean.

B) Venn diagram showing overlap between the top 25% Pol II-occupied genes and those genes which were identified to express cryptic transcripts in *spt16-197* and *spt6-1004* (Cheung et al. 2008).

C) Overlapping genes, identified to express cryptic transcripts and were among the top 25% Pol II occupied genes (n=154), were removed from the list of 1246 highly expressed genes. Pol II occupancy profile for the remaining 1092 genes in WT, spt16 and spt6 cells is shown.

D) Rpb3 occupancy profile of WT and *spt6* cells at the genes eliciting reduction in Rpb3 occupancy ≥ 0.5 log_2_ ratio (ChIP/input) upon depleting Spt6.

E) The top 25% Pol II-occupied genes were grouped on the basis of their gene-length and changes in Rpb3 occupancy profiles (*spt6*/WT) for the long (>2 kb), medium (1-2 kb), and short (0.5-1 kb) are shown in the WT and Spt6-depleted cells.

**Figure S4: Gene-length analyses for the genes most affected by depletion of Spt6 or Spt16.**

A) The genes which showed more than 0.5 log_2_ reduction in Pol II occupancy upon depleting either Spt6 or Spt16 were analyzed, and were classified as Spt16-unique, Spt6-unique and Spt16-Spt6-common depending on whether they were enriched only upon depleting Spt16, Spt6 or both, respectively. Frequency distribution of these classes of genes based on their gene-length is plotted as a histogram.

B) Gene-length for the three classes of genes is shown in the box plot. Center lines show the medians; box limits indicate the 25th and 75th percentiles as determined by R software; whiskers extend 1.5 times the interquartile range from the 25th and 75th percentiles, outliers are represented by dots. 115, 217 and 111 genes were present in the Spt16-unique, Spt6-unique and common gene-sets, respectively.

**Figure S5: Pol II occupancy in *spt16/spt6*Δ*202* cells.**

A) Western blot showing Spt16 protein levels in untreated and dox-treated *Spt16/spt6*Δ*202* cells.

B) Rpb3 occupancy profiles for the top 25% Pol II-occupied genes (n=1246) in untreated *SPT16/spt6*Δ*202* and dox-treated (*spt16/spt6*Δ*202*) cells.

C) Box-plot showing Rpb3 enrichment (log_2_ ratio, ChIP/input) for the 137 highly expressed genes among the 200 genes showing greatest reduction in *Spt16/spt6*Δ*202* dox-treated compared to untreated cells. Pol II occupancies are shown for the untreated WT and *SPT16/spt6*Δ*202*, and dox-treated *SPT16-TET (spt16*) and *SPT6-TET (spt6*) and *SPT16/spt6*Δ*202 (spt16/spt6*Δ*202*) cells.

